# Reduced carbohydrate complexity alters gut microbial structure independent of total carbohydrate intake

**DOI:** 10.1101/2025.09.20.677466

**Authors:** Clara Flores, Anna M. Seekatz

## Abstract

Dietary habits have dramatically altered over recent decades, yet the impact of simplified carbohydrate intake on the gut microbiome’s complexity and function remains poorly understood. This study investigates how the variety of dietary carbohydrates—not just their amount—shapes gut microbial diversity and resilience in C57BL/6 mice. Over eight weeks, mice consumed diets varying in carbohydrate complexity but matched for total carbohydrate content. Using 16S rRNA sequencing, we found that reduced carbohydrate diversity led to significant declines in microbial diversity and taxonomic redundancy among important bacterial groups, such as unclassified *Lachnospiraceae, Ruminococcaceae*, and *Muribaculaceae*, despite no immediate changes in host physiology. Concurrently, *Akkermansia* increased under low-complexity diets, suggesting a shift toward mucin degradation when complex polysaccharides are scarce. These changes indicate that loss of carbohydrate complexity narrows microbial niches, potentially disrupting metabolic interactions and functional stability of the gut ecosystem. Given the widespread adoption of processed, low-fiber diets in modern societies, these findings emphasize the importance of macronutrient complexity in maintaining gut microbial health. While short-term host effects were minimal, the microbial shifts observed could presage long-term consequences for gut resilience and disease susceptibility. This study underscores the need to consider carbohydrate diversity in dietary recommendations and microbial ecology research to safeguard gut health in the face of global dietary simplification.

## INTRODUCTION

The gut microbiome, estimated to harbor microbes that encode up to 100-fold more unique genes than our own genomes, plays a vital role in supporting host functions such as immune system development (Qin et al., 2010), colonization resistance against pathogens (I. Khan et al., 2021), and the production of necessary metabolites (Jyoti & Dey, 2025). Despite millennia of mutualistic coevolution between hosts and microbes that have established discrete microbiomes to each of their hosts (Dethlefsen et al., 2007; Koskella & Bergelson, 2020), this environment is dynamic and susceptible to microbial shifts in response to environmental triggers. Physical exercise, xenobiotics, antibiotics, and diet are among some of the factors that have been linked to changes in the gut microbiome (Gacesa et al., 2022), with diet being one of the most modifiable determinants of microbial community composition (Sanz et al., 2025). Both short and long-term dietary patterns have been demonstrated to induce measurable changes in microbial taxa and metabolic outputs (Grembi et al., 2020). Moreover, dietary interventions have been associated with clinically relevant improvements in chronic disease outcomes, including metabolic syndrome, inflammatory bowel disease, and type 2 diabetes (Gropper, 2023). Given the central role of diet in shaping gut microbiota, investigating how changes in diet influence microbial composition are key to understanding its impact on health and disease.

Diet is a major contributor to the nutrient landscape of the gut, shaping the availability of metabolic substrates such as carbohydrates and amino acids that influence the taxonomic distribution and diversity of gut microbes (Leeming et al., 2019; Zhang, 2022). Long-term diet studies have shown an increase in *Bacteroides* for healthy human volunteers that consumed higher levels of protein and animal fat, while those consuming mostly carbohydrates had higher rates of *Prevotella* (Wu et al., 2011). Additionally, studies in mice have demonstrated that even within a short 2-week period, feeding mice a high-protein/low-carbohydrate diet decreased the proportion of Lachnospiraceae and Ruminococcaceae in the gut and increased proportions of *Bacteroides* and *Parabacteroides* compared to mice fed a normal protein/carbohydrate diet (Kim et al., 2016). Moreover, in a study in humans, a 2-week dietary fiber intervention resulted in increased *Bifidobacterium* and *Lactobacillus* (A. Oliver et al., 2021). Dietary influences on taxonomic ratios in the gut can, in turn, influence health and disease. For instance, decreased abundance of Lachnospiraceae, key butyrate producers in the gut, have been correlated with higher blood pressure, total cholesterol and triglycerides (Ahrens et al., 2021). Increased abundance of *Parabacteroides* has been reported to be correlated with hypertension, although its presence has also been associated with regulating immunity, relieving inflammation, and carbon metabolism (Cui et al., 2022). While these findings highlight the microbiome’s responsiveness to dietary inputs, our understanding of exactly how macronutrient complexity change resources available to specific gut members, and thus shapes community composition, remains incomplete.

Of the macronutrients our diets supply, carbohydrates, both simple and complex, serve as a primary energy source for humans and their associated gut microbes (Mora-Flores et al., 2023). While monosaccharides and disaccharides are rapidly absorbed in the small intestine, complex carbohydrates, particularly dietary fibers and resistant starches, escape host digestion and reach the colon where they are fermented by diverse bacterial populations. This fermentation process produces key microbial metabolites, including short-chain fatty acids (SCFAs), tryptophan-derived indoles, and secondary bile acids, which are known to support gut health and function (Postler & Ghosh, 2017). Importantly, different microbial taxa exhibit specialized enzymatic capacities that enable them to degrade complex carbohydrates, largely through the expression of glycoside hydrolases (GHs). These enzymes cleave glycosidic bonds in polysaccharides and oligosaccharides, and their diversity within microbial genomes reflects taxon-specific metabolic niches (Berkhout et al., 2022; Takihara & Okuda, 2023). The structural complexity and heterogeneity of dietary carbohydrates, therefore, serve as ecological filters that shape microbial community composition by selecting for microbes capable of degrading specific glycan structures. Diets rich in a variety of complex carbohydrates can promote greater taxonomic diversity by supporting a broader array of glycan utilizing microbes, whereas simplified carbohydrate inputs may promote narrower microbial niches leading to reduced diversity and potential functional loss (Takihara & Okuda, 2023).

Historically, human diets were rich in complex carbohydrates, including fiber, which has well-documented benefits for host health (Alcock et al., 2021). With urbanization and the rise of modern agricultural and food processing practices, much of Westernized diets have decreased in complex carbohydrate consumption compared to ancestral diets (Casari et al., 2022). Modern, Western diets are characterized by foods high in fat, sugar, and salt, pre-packaged or processed foods, and low in fiber (Clemente-Suárez et al., 2023). Studies show that such diets have been associated with increased allergies, asthma, chronic autoimmune diseases, inflammation, reduction in species richness in both Bacteroidetes and Firmicutes (Huang et al., 2023) and bacterial blooms in Mollicutes in mice which have been implicated in increased host adiposity (Turnbaugh et al., 2008). While these compositional shifts are significant, the ecological consequences extend further, affecting key ecosystem traits including species and functional diversity, stability, and taxonomic and functional redundancy. Taxonomic diversity supports ecosystem stability by maintaining functional redundancy, whereby multiple microbial species can perform similar metabolic roles (Eisenhauer et al., 2023). Conversely, reduced diversity, often observed in response to Western dietary patterns, can lead to reduced taxonomic redundancy –similar or closely related species—which may signal a decline in functional redundancy leaving ecosystems vulnerable to functional losses after disturbance (Cross et al., 2025; T. H. Oliver et al., 2015; Trivedi et al., 2019). These shifts can drive the microbiome into alternate stable states that are less favorable to host health (Flores et al., 2025; Van de Guchte et al., 2020). Therefore, understanding how specific dietary features, particularly the complexity of carbohydrate inputs, influence microbial diversity and redundancy is essential for elucidating mechanisms that underlie the link between modern diets and gut microbial structure. This raises a key question: what happens to microbial communities when complex nutrient sources are simplified?

In this study, we aimed to investigate how digestible carbohydrate complexity influenced taxonomic diversity and redundancy in the murine gut microbiome. Mice were fed customized diets that varied in the number of different types of carbohydrates, rather than amount of total carbohydrates, for a total of eight weeks. Using 16S rRNA gene-based sequencing, we compared phylogeny, taxonomic diversity and redundancy within and between our three groups of mice. We observed decreased overall taxonomic diversity and redundancy for select taxa in mice fed mid and low complexity diets in comparison to mice fed a high complexity diet. While taxonomic redundancy was reduced in some genera, including *Lachnospiraceae* and *Muribaculaceae*, other genera, including *Akkermansia*, maintained similar levels of taxonomic redundancy, suggesting that the observed shifts in the gut microbiota were not universally distributed across all taxa and overall diversity.

## MATERIALS AND METHODS

### Animals and diets

C57BL/6 mice (6 to 10 weeks old) were purchased through Jackson Laboratory. All experiments using mice were conducted under protocols approved by Clemson University’s Institutional Animal Care and Use Committee. Upon arrival mice were housed, by sex, randomly assigned to groups of three per cage under the same conditions on a 12 h light/dark cycle. All cages, bedding (7084 Teklad Pelleted Paper Bedding), and water were autoclaved, and mice were given γ-irradiated custom diets from Research Diets, New Brunswick, NJ (Table S1). For the first 4 weeks mice were fed *ad libitum* a high complexity diet (HCD, 50% complex carbohydrates from corn starch, wheat, and potato starch and 50% simple carbohydrates; D22021105). After the 4-week acclimatation period, mice were either maintained on the HCD (n = 18) or fed a mid-complexity diet (n = 18; MCD, 50% complex carbohydrates from corn starch, and 50% simple carbohydrates; #D22021106) or low complexity diet (n = 18; LCD, 87.5% simple carbohydrates; #D22021107). All three diets were tested on equal numbers of male and female mice in two separate experimental replicates (Table S2).

### Intestinal barrier integrity and gastrointestinal transit time

To assess intestinal barrier function, mice were fasted for three hours with *ad libitum* access to water prior to oral gavage with 0.2□mL of sterile saline containing 12□mg fluorescein isothiocyanate (FITC)-dextran (4□kDa), 8□mg rhodamine B (70□kDa), and 20□mg creatinine (all reagents from Sigma-Aldrich; #46944, #R9379, and #C4255). After five hours, blood was collected from anesthetized mice via retro-orbital bleed, and serum was isolated by centrifugation. Fluorescence in serum samples was quantified using a Synergy HT plate reader (BioTek) against freshly prepared standard curves. FITC was measured with excitation/emission wavelengths of 495/525□nm, and rhodamine B was measured at 555/585□nm, respectively.

Gastrointestinal transit time was assessed by oral gavage of 150□µL of a 6% (w/v) carmine red dye solution prepared in 0.5% methylcellulose. Mice were fasted from both food and water for the duration of the assay. Following gavage, mice were individually housed in sterile 1□mL pipette boxes lined with an absorbent pad. Fecal pellets were examined every 15 minutes, and transit time was defined as the interval between gavage and first appearance of red-colored feces. Mice were returned to their home cages immediately after dye detection.

### Dual-energy X-ray absorptiometry (DXA) scan

Animals were weighed prior to euthanasia, after which cecal contents and tissue, ileal contents, and colonic contents and tissue were collected. Carcasses were kept on ice until transport to the dual-energy X-ray absorptiometry (DXA) facility. Upon arrival, animals were allowed to thaw at room temperature and were positioned in a standardized left lateral recumbent orientation for scanning. Whole-body composition was assessed using a Hologic DXA scanner (Discovery A, serial number 87637), operated using the manufacturer’s rat whole-body scan settings. Scans were analyzed using the instrument’s standard analysis software, and the following parameters were recorded: total area, bone mineral content (BMC), bone mineral density (BMD), fat mass, lean mass plus BMC, total mass, and percent body fat.

### 16S rRNA gene-based sequencing

DNA isolation was done on mouse fecal and cecal content using Qiagen’s MagAttract PowerMicrobiome DNA/RNA kit (Cat No. 27500-4-EP). The bacterial V4 16S rRNA region was amplified as described previously (J. J. Kozich et al., 2013). Briefly, amplicons were generated using a high-fidelity polymerase (Invitrogen, #12-346-086). Libraries were cleaned up and normalized using Invitrogen SequalPrep Plate Normalization kit (#A1051001). After normalization, sample quality was assessed using the Agilent TapeStation and quantified using KAPA biosystems qPCR kit. Samples were sequenced on Illumina’s MiSeq platform following the Schloss lab MiSeq Wet Lab standard operating procedure (SOP) with a 10% PhiX spike according to the manufacturer’s protocol using a final concentration of 4pM (J. Kozich et al., 2013; J. J. Kozich et al., 2013). Raw sequence data has been deposited in the Short Read Archive, under BioProject PRJNA1308500 (SAMN# 50775611 – 50776048).

### Microbiome analysis

For 16S rRNA gene-based analysis, sequences were processed using mothur v.1.48.0 (J. J. Kozich et al., 2013). Briefly, the SILVA rRNA database project (v132) (Quast et al., 2012) was used to align reads to the v4 region of the 16S rRNA gene. Sequences were clustered by 100% sequence similarity into amplicon sequence variants (ASVs) and classified using the mothur-adapted version of the Ribosomal DNA Project (RDP) database (v18) (Cole et al., 2014). All statistical analyses and data visualization were performed using R (v4.4.0) with dplyr, readaxl, and ggplot2 (Wickham et al., 2025; Wickham & Bryan, 2025; Wickham, 2016) used for data visualization. Principal coordinates analysis (PCoA) and permutational multivariate analysis of variance (PERMANOVA) were conducted using the vegan package (Oksanen et al., 2025). Data distributions were assessed using the Shapiro-Wilk test for normality and Levene’s test for homogeneity of variance. For comparisons across multiple groups, the Kruskal–Wallis test was used, followed by post hoc pairwise Dunn’s test or Wilcoxon rank-sum tests where appropriate. Differentially abundant ASVs between lower complexity diets and HCD-fed mice were identified using Multivariable Association with Linear Models (MaAsLin2) (Mallick et al., 2021). All specific commands and quality parameters are available at https://github.com/SeekatzLab/RoL-FunctionalRedundancy-microbiota.

## RESULTS

### Diets varying in carbohydrate complexity did not impact host health

To test the effects of dietary carbohydrate complexity on the diversity and taxonomic redundancy of the gut microbiota, we developed three custom diets that exhibited the same percentage of total carbohydrates but differed in composition and number of carbohydrates (Figure 1A, Table S1). C57BL/6J mice (n = 54) were all maintained on the high complexity diet (HCD) for four weeks, then either continued on the HCD or switched to a mid-complexity (MCD) or low complexity diet (LCD) (n = 18 per group; Figure 1B). Mice in all groups steadily gained weight throughout the study (Figure 1C) and although total weight gain did not differ significantly across diet groups (Figure 1D), the rate of weight gain for mice fed LCD was significantly slower compared to those maintained on HCD (linear mixed-effects model, p < 0.0001). Though female mice consumed fewer weekly and total kilocalories (kcals) than male mice, mice fed lower complexity diets did not consume significantly more total kcals than mice maintained on HCD (Figure S1A-D). Moreover, a subset of mice (n = 18; 6 per group) scanned for body mass composition metrics, including total fat mass, using dual-energy X-ray absorptiometry (DEXA) did not exhibit differences between treatment groups (Figure S1B-G). Gastrointestinal (GI) transit time (Figure S2A) and intestinal barrier integrity (Figure S2B-C) were also not significantly impacted by any of the diet switches.

**Figure 1.**
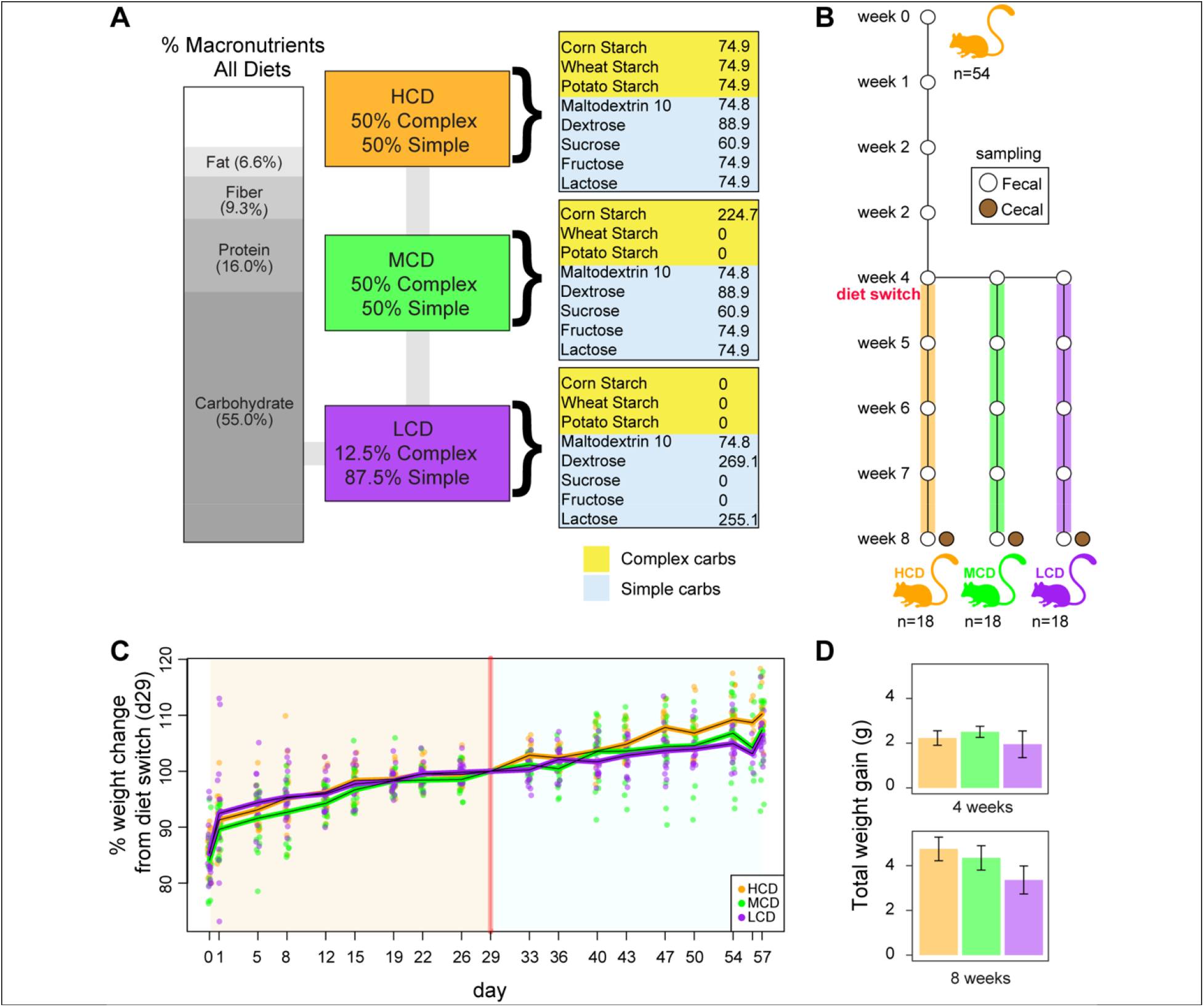
Experimental design and dietary intervention. **A)** Composition of customized diets designed to measure shifts in gut microbiota taxonomic redundancy in response to carbohydrate complexity. All diets were isocaloric and matched for total macronutrient content (fat, protein, carbohydrate, and fiber), varying only in composition of carbohydrate sources (proportions of complex vs. simple sugars). **B)** Schematic of experimental design using a mouse model. **C)** Body weight trajectories of mice over the 8-week study period. Weights were normalized to the day of diet switch (d29). Left panel shows weight change during the initial four weeks (pre-diet switch), right panel shows weight change during the final four weeks (post-diet switch). **D)** Total weight gain (g) per diet group, averaged over the first four weeks and over the full 8-week study period.

### Lower complexity diets decreased microbiota diversity and shifted community composition

We next sought to identify how the diets impacted the gut microbiota using 16S rRNA gene-based sequencing. Mice fed diets of decreasing complexity (MCD and LCD) demonstrated an overall decrease in diversity in their fecal microbiota following the diet switch and in cecal microbiota at the conclusion of the study, as measured by the inverse Simpson index (Figure 2A, B). Similar patterns were observed across additional alpha diversity metrics analyzed (Figure S3A-D), with Wilcoxon rank-sum pairwise comparisons indicating significant differences (p < 0.05). Changes to the microbiota structure recapitulated the observed changes in diversity, as samples collected from mice fed MCD or LCD clustered away from mice fed HCD or samples collected prior to the diet switch, based on the Bray-Curtis dissimilarity calculated from amplicon sequence variants (ASVs) (Figure 2C, AMOVA, p < 0.005). A biplot based on cecal ASVs overlayed over the observed microbiota structural comparisons revealed four ASVs driving community structure, with ASV01 (*Akkermansia)* driving MCD and LCD group clustering, and ASV02 (unclassified *Muribaculaceae species)*, ASV04 *(Dubosiella)*, and ASV06 (unclassified *Lachnospiraceae* species) driving the pre-diet change and HCD community clustering (Figure 2C). This separation was also evident in endpoint cecal samples, where mice fed HCD were more dissimilar from those fed either MCD or LCD (Figure 2D). Similarly, while the fecal microbiota of mice maintained on HCD did not differ significantly from their baseline samples, mice fed either MCD or LCD demonstrated a rapid shift in community structure after the diet switch (Figure 2E)

**Figure 2.**
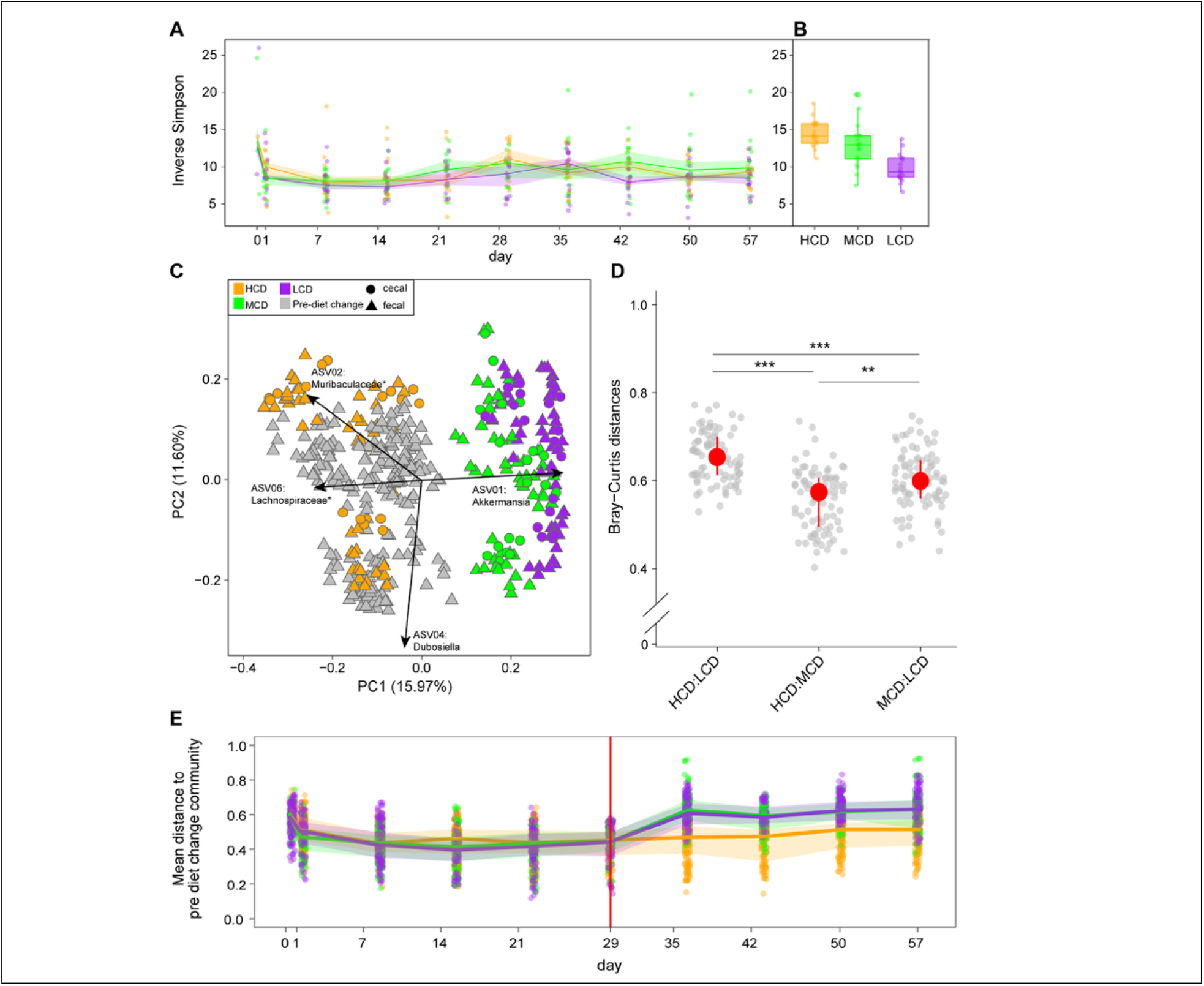
Mid and low complexity diets reduce taxonomic diversity. **A-B)** Inverse Simpson index of **A)** fecal samples collected throughout the study and **B)** cecal samples collected at the end of the study. **C)** Principal Coordinates Analysis of Bray-Curtis dissimilarity based on ASV abundances from fecal and cecal samples (AMOVA, *p < 0.001). Arrows represent ASVs contributing to the clustering of cecal communities by diet. **D)** Pairwise Bray-Curtis dissimilarity between cecal samples by diet group. **E)** Bray-Curtis dissimilarity over time, calculated as distance from the pre-diet switch (d29). Statistical significance for panels A, B, D, and E assessed using Kruskal-Wallis with post hoc Dunn’s test (*p<0.05, **p<0.001, ***p<0.0001).

### Lower-complexity diets increase *Akkermansia* and decrease ASVs belonging to the family Lachnospiraceae

Taxonomic changes in the microbiota mirrored diversity and structural changes due to diet. Across all groups, taxonomic profiling identified genera belonging to six phyla, Actinomycetota (formerly Actinobacteria), Bacillota (Firmicutes), Bacteroidota (Bacteroidetes), Pseudomonadota (Proteobacteria), Mycoplasmatota (Tenericutes), and Verrucomicrobiota (Verrucomicrobia), plus unclassified bacteria. Together, Bacillota and Bacteroidota constituted the majority of the phyla observed across all groups of mice (Figure 1A), consistent with a healthy murine microbiota. However, within the cecal microbiota, HCD-fed mice exhibited a significantly lower Bacillota/Bacteroidota ratio compared to MCD- and LCD-fed mice (mean ratios: HCD = 2.9; MCD = 4.24; LCD = 4.23; Wilcoxon, p < 0.005) (Figure S3E).

Verrucomicrobiota and Mycoplasmatota were each represented by multiple ASVs representing a single genus, *Akkermansia* and *Anaeroplasma*, respectively.

Significantly higher abundances of *Akkermansia*, mainly contributed to by ASV01, were observed in MCD- and LCD-fed mice (average 19.1% and 22.4%) compared to HCD mice (9.5%) (Figure 3A, Wilcoxon, p < 0.001). In contrast, *Anaeroplasma*, primarily contributed to by ASV034, was decreased in mice fed MCD and LCD (0.5% and 0.02%) compared to HCD-fed mice (1.6%). Some experimental variation was observed across cage replicates and/or sex within the groups, most notably by the presence of increased Actinomycetota in cages 907-910 and increased Pseudomonadota in cages 911-912 and 915-916, which represented a second experimental validation (Table S2). In these mice, Actinomycetota, primarily composed of *Bifidobacterium* (ASV003), was significantly decreased in MCD and LCD mice (average 0.05% and 0.25%) compared to HCD mice (9.8%) (Wilcoxon, p < 0.001). While only contributing a small percentage of the total relative abundance, *Parasutterella*, belonging to the phylum Pseudomonadota, was increased in these MCD and LCD mice (0.59% and 0.42%) compared to HCD mice (0.23%), the effects of which were enhanced in female compared to male mice (Figure 3A). These findings corroborated the previously observed correlating ASVs in our biplot results (Figure 2B).

**Figure 3.**
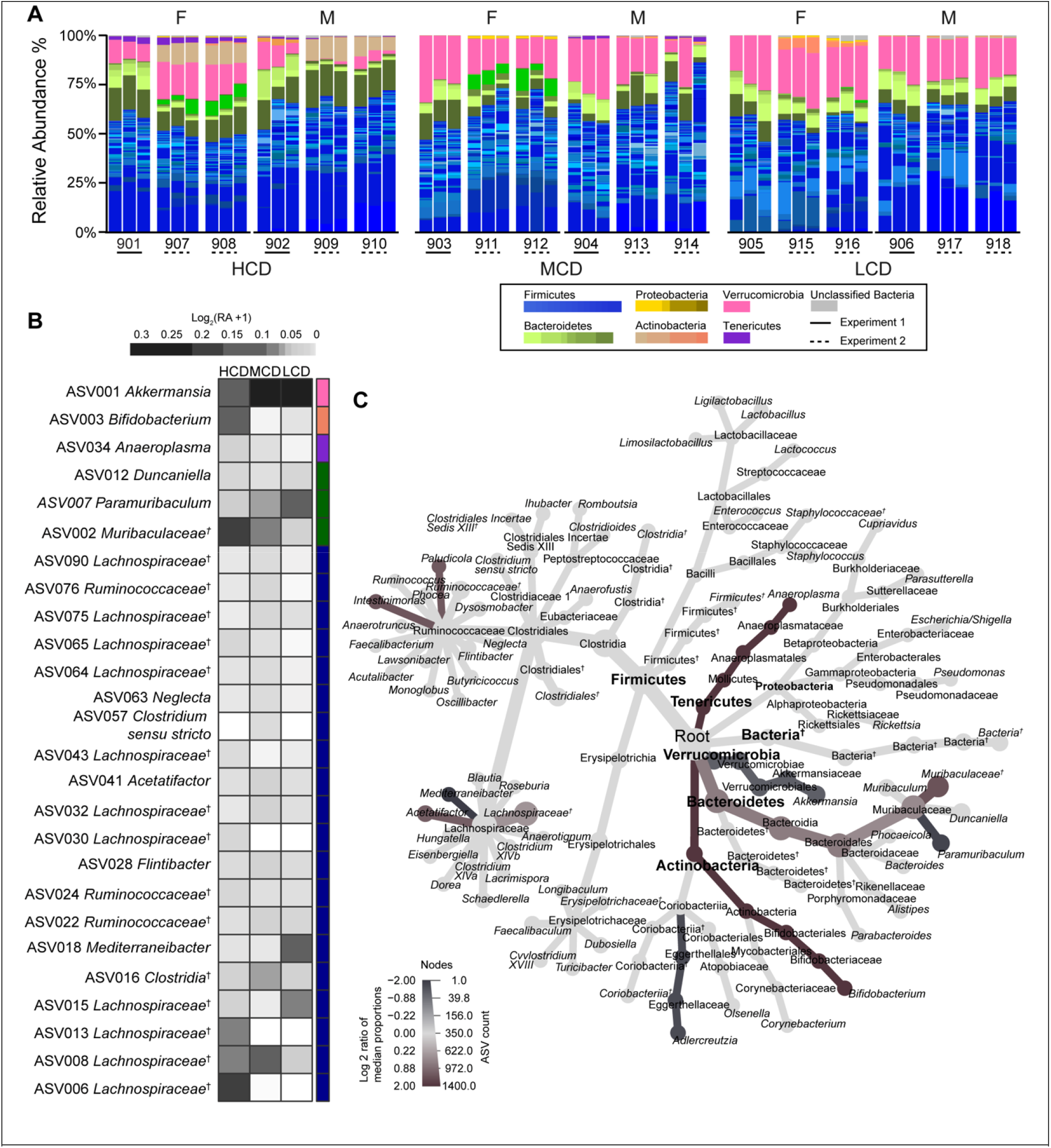
Lower-complexity diets increase *Akkermansia* and decrease unclassified *Lachnospiraceae*. **A)** Relative abundance of cecal ASVs grouped by diet (HCD, MCD, and LCD), cage ID (901 – 918), and sex (F = female; M = male). Legend color-coded by phylum/genus. **B)** Mean logD-transformed relative abundance (RA + 1) of genera identified as significantly different across diet groups compared to HCD-fed mice (MaAsLin2; linear model with Benjamini-Hochberg correction, q ≤ 0.005). Right-side color bar corresponds to phylum-level legend in panel A. **C)** Heat tree comparing LCD- and HCD-fed mice, showing the log□ ratio of median taxon proportions. Node size represents the number of samples containing each taxon; color denotes the number of ASVs assigned to each taxon. Statistical significance assessed using Wilcoxon rank-sum test with FDR correction. ^†^ Denotes unclassified taxa.

We next assessed which of these taxa were differentially abundant in LCD and MCD groups compared to HCD-fed mice using MaAsLin2, filtering for a minimum relative abundance of at least 0.001 for each ASV. Across experiments, *Akkermansia* (ASV001) was significantly increased in mice fed MCD and LCD, with nearly twice the overall relative abundance across all mice and near absence in male mice in cage 909 fed HCD (Figure 3B). Among the top 26 differentially abundant ASVs, 11 were unclassified *Lachnospiraceae*, not all of which exhibited the same pattern of differential abundance across the diets. For instance, unclassified *Lachnospiraceae* ASVs 006, 013, and 030 were increased in HCD-fed mice, while ASV008 was increased in MCD-fed mice, and ASV015 was increased in LCD-fed mice.

Other taxa significantly decreased in mice fed diets of lower complexity included *Bifidobacterium (*ASV003) and unclassified *Muribaculaceae* (ASV002) (Figure 3B). Given that the HCD and LCD represented the most contrasting dietary treatments in terms of carbohydrate complexity and diversity and exhibited the largest shifts in community composition and structure, we next conducted a focused comparison between these two groups to further contextualize the observed taxonomic differences. While MaAsLin2 highlighted individual ASVs that were significantly altered between the lower complexity diets and HCD-fed mice, we used a complementary taxon-level approach to explore broader shifts in community structure between HCD- and LCD-fed mice. This analysis revealed distinct patterns of taxon richness and prevalence, with *Anaeroplasma, Bifidobacterium*, unclassified *Muribaculaceae*, and *Intestimonas* more commonly detected in HCD-fed mice, whereas *Akkermansia, Paramuribaculum*, and *Adlercreutzia* were more prevalent in LCD-fed mice (Figure 3C). Within the Lachnospiraceae family, which had one of the highest numbers of assigned ASVs, a majority of the unclassified *Lachnospiraceae* and *Acetatifactor* ASVs were significantly more prevalent in HCD-fed mice while *Mediterraneibacter* was more prevalent in LCD-fed mice (Figure 3C).

### Mice fed lower complexity diets exhibited lower taxonomic redundancy for select taxa

To better understand how the customized diets shaped the gut microbiota, we quantified taxonomic redundancy by determining, at each taxonomic rank (phylum to genus), the number of unique ASVs assigned to each taxon. This allowed us to assess how ASV-level diversity was partitioned across broader taxonomic categories, providing insight into how microbial diversity is organized within different taxonomic levels. At the phylum level, we observed increased Bacillota, Bacteroidota, and Mycoplasmatota unique ASVs in mice fed HCD (Figure 4A). At the genus level (Figure 4B), the top three genera with the highest ASV counts (unclassified *Lachnospiraceae*, unclassified *Ruminococcaceae*, and unclassified *Clostridiales*) belonged to the Bacillota phylum. Among these, unclassified *Lachnospiraceae* showed the highest taxonomic redundancy in HCD-fed mice (average 67.4), significantly more than in MCD (62.7) and LCD (45.4) groups (Wilcoxon, p < 0.05).

**Figure 4.**
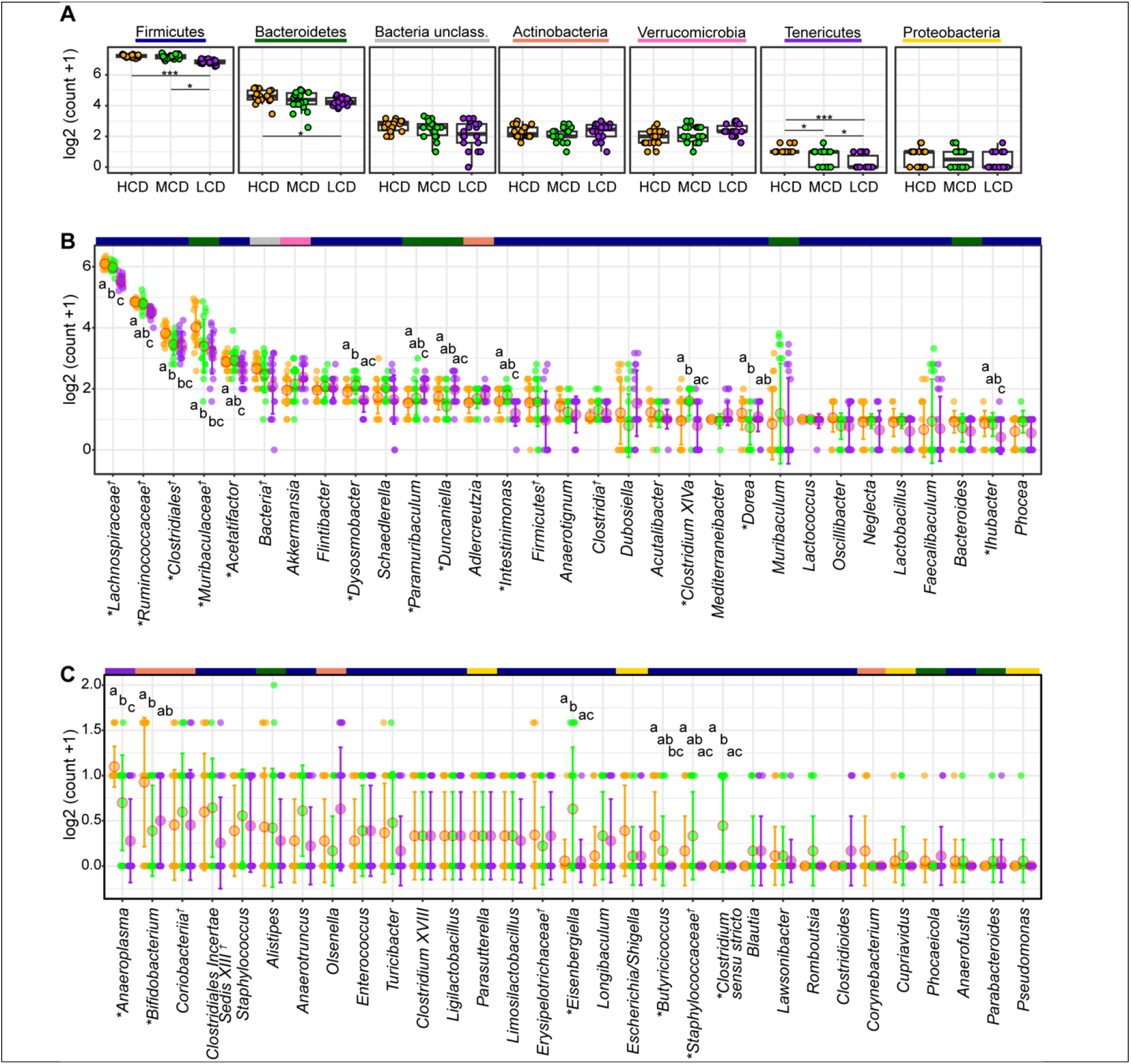
Low-complexity diets resulted in reduced taxonomic redundancy for ASVs observed in *Lachnospiraceae* but not *Akkermansia*. **A)** log□ (count + 1) of unique ASV counts per mouse at the phylum level. **B–C)** log□ (count + 1) of unique ASV counts per genus for the top half **B)** and bottom half **C)** of genera, ranked by mean ASV count across all samples. Color-bars correspond to phylum-level classifications of displayed genera, as displayed in panel A phylum groups. Statistical comparisons for all analyses performed using Kruskal-Wallis test with post hoc Wilcoxon rank-sum tests (*p<0.05, **p<0.001, ***p<0.0001). * Indicate significant differences between at least one pairwise group comparison. ^†^ Denotes unclassified taxa.

Taxonomic redundancy decreased in mice fed LCD within the unclassified *Ruminococcacea*, unclassified *Clostridiales*, unclassified *Muribaculaceae*, and unclassified *Acetatifactor* genera, mirroring the general pattern of decreased relative abundance in these mice (Figure 4B). While Verrucomicrobiota, represented by a single genus (*Akkermansia)*, demonstrated no significant changes across taxonomic levels, Mycoplasmatota, also represented by a single genus (*Aneroplasma*), had significantly increased redundancy in HCD-fed mice at all taxonomic levels (Figures S4-S6). Moreover, mice fed LCD had significantly higher ASV counts for *Paramuribaculum* and *Duncaniella*, both of which belong to the phyla Bacteroidota.

Specifically, *Duncaniella* was significantly more taxonomically redundant in LCD-fed mice compared to MCD-fed mice (Figure 4B).

While most genera did not exhibit statistically significant differences in taxonomic redundancy across diet groups, several did—most of which belonged to the phylum Bacillota (Figure 4B-C, Table S3). Non-significant but consistent trends in taxonomic redundancy were observed from phylum to genus for unclassified bacteria, Pseudomonadota, and Verrucomicrobiota. These patterns are illustrated at the class (Figure S4), order (Figures S5), and family (Figure S6) levels. In contrast, Actinomycetota deviated from the taxonomic redundancy patterns observed at the phylum level. At the family level, Bifidobacteriaceae (Figure S6) was significantly more taxonomically redundant in HCD-fed mice. Notably, this taxon was absent in the first experiment, highlighting inherent variability of the microbiota between experimental cohorts.

## DISCUSSION

In this study, we demonstrate that dietary carbohydrate complexity, independent of total carbohydrate content, influences the diversity, composition, and taxonomic redundancy of the gut microbiota in C57BL/6 mice, with minimal measured effects on host physiology. These findings provide evidence that macronutrient composition—not just quantity—is a critical determinant of gut microbiota structure and function. Reduced carbohydrate complexity was associated with a significant decrease in *Lachnospiraceae* species and a corresponding increase in *Akkermansia*, driving shifts in microbial community structure of MCD- and LCD-fed mice away from their pre-diet change and HCD-fed counterparts. Although these and other taxa were differentially abundant in MCD and LCD groups, this did not always correlate with a decrease in taxonomic redundancy, albeit redundancy did decline for select taxa including *Lachnospiraceae, Ruminococcaceae* and *Muribaculaceae* in LCD fed mice. These findings, combined with existing research linking microbiome composition to inflammation and gastrointestinal disease (Elmassry et al., 2025; Madhogaria et al., 2022; Schirmer et al., 2018; Turnbaugh et al., 2006; Zhao et al., 2023), underscore the need to more thoroughly investigate dietary carbohydrate complexity as a critical modulator of gut microbial ecology and host health.

Despite equivalent macronutrient caloric content across all diets, we observed a clear reduction in diversity and taxonomic redundancy for select taxa as carbohydrate complexity decreased. These patterns mirror previous findings that microbiota composition and structure are particularly sensitive to sugar composition and fiber content (Beilharz, Kaakoush, et al., 2016; Faits et al., 2020; S. Khan et al., 2020; Laffin et al., 2019; Menni et al., 2017). Gut microbiota appeared highly responsive to even subtle alterations in substrate complexity, as evidenced by distinct clustering of LCD- and MCD-fed mice from those consuming HCD. In this context, differences in starch composition likely played a role in shaping microbial community structure. The HCD contained corn, wheat and potato starch, a known source of resistant starch (Bang et al., 2019; Romano et al., 2016; Singh et al., 2003), which resists digestion in the small intestine and undergoes fermentation in the colon, effectively extending the pool of fermentable substrates for saccharolytic taxa. In contrast, the MCD included corn starch as its sole starch source, while the LCD was devoid of complex starches entirely. This progressive narrowing of fermentable substrate availability may have imposed niche restrictions, particularly for specialist taxa able to use complex polysaccharides for growth. Resistant starches not only support taxa directly involved in their fermentation, such as *Parabacteroides, Ruminococcus*, and *Bifidobacterium*, but also facilitate the expansion of secondary fermenters through cross-feeding interactions (Dobranowski & Stintzi, 2021; Jung & Park, 2023; Martínez et al., 2010). For instance, increases in taxa not directly involved in resistant starch degradation have been observed in resistant starch-fed animals (Hu et al., 2016), likely due to their utilization of fermentation byproducts such as oligosaccharides, SCFAs, or lactate (Dobranowski & Stintzi, 2021). These metabolic interactions help sustain trophic networks, maintain redox balance, and promote beneficial functions such as mucosal regeneration and inflammation control (Shi et al., 2017; Valiei et al., 2024). Therefore, the loss of complex carbohydrates may disrupt these interdependent relationships, limiting metabolic flexibility, constraining the functional potential of the gut microbiota and ultimately diminish community resilience.

Notably, the substantial microbial restructuring we observed occurred without changes in body weight, suggesting a degree of short-term host resilience to microbial shifts. This is consistent with Do et al., who observed high-glucose and high-fructose diets led to significant alterations in gut microbiota composition without accompanying elevations in body weight (Do et al., 2018). However, they did report elevated blood glucose and endotoxin levels, indicating early signs of metabolic disruption. In line with our findings, other rodent studies employing higher-sugar diets have similarly reported no significant changes in body weight or total caloric intake, despite the presence of physiological or metabolic perturbations (Beilharz et al., 2014; Beilharz, Maniam, et al., 2016). Moreover, others have observed a diet rich in simple sugars promotes a pro-inflammatory response via gut microbiota shifts (Fajstova et al., 2020), further supporting the notion that microbial alterations may precede overt metabolic dysfunction. Taken together, these compositional shifts could reflect the narrowing of available metabolic niches under simplified carbohydrate conditions and highlight the sensitivity of gut microbial ecosystems to reductions in dietary complexity, an increasingly relevant concern in modern diets characterized by highly homogenized, processed foods (Clemente-Suárez et al., 2023).

One of the most striking taxonomic changes was the increase in relative abundance of *Akkermansia* in mice fed lower-complexity diets accompanied by a reduction of *Lachnospiraceae* species. *Akkermansia* was also differentially abundant in MCD- and LCD-fed mice compared to HCD fed mice, echoing findings from Khan et al (2020), where diets supplemented with 10% glucose similarly led to an enrichment of Akkermansiaceae and a decline of Lachnospiraceae (S. Khan et al., 2020). In contrast to fructose and sucrose simple sugar supplementation used in many studies (Fisher et al., 2024; Sun et al., 2021), our LCD formulation contained a combination of dextrose and lactose alongside lower levels of more complex carbohydrates. This suggests that the observed *Akkermansia* enrichment may reflect compensatory mucin utilization in the relative absence of diverse dietary polysaccharides, a hypothesis supported by prior studies showing increased *Akkermansia* abundance under fiber- or polysaccharide-deprived conditions (Desai et al., 2016; Wolter et al., 2024). Limiting nutrient diversity can lead to increased microbial competition, which often results in reduced community diversity as dominant species outcompete others for scarce resources (Brochet et al., 2021; Yu et al., 2024).

Though taxonomic redundancy remained largely unchanged across diet groups, MCD- and LCD-fed mice exhibited significantly decreased redundancy for many Bacillota, including, *Clostridiales, Dysomobacter*, and *Lachnospiraceae*, a group broadly associated with health benefits, largely due to their capacity to produce SCFAs (Vacca et al., 2020). Lachnospiraceae, a taxonomically and functionally diverse family composed of both generalists and specialists (Sorbara et al., 2020) have been shown to suppress inflammation, and their reduced abundance has been linked to increased susceptibility to colitis (Frank et al., 2007; Vacca et al., 2020).

Some members may be better adapted to flexible nutrient environments, while others rely more heavily on specific substrates that may include complex carbohydrates whose hydrolysis depends on available glycoside hydrolases (GHs) (Flint et al., 2012). In the absence of exogenous carbohydrate sources, as in the MCD and LCD diets, specialist taxa within Lachnospiraceae may become competitively disadvantaged. In contrast, *Akkermansia muciniphila* is specifically suited to use host mucins as nitrogen and carbon sources (Yamaguchi & Yamamoto, 2023). This specialization sets *Akkermansia* apart from other generalist microbes, which are not adapted to utilize host-derived glycans (Berkhout et al., 2022; Kostopoulos et al., 2021). The differences in GH repertoire may confer a competitive edge to mucin-degrading Verrucomicrobia under carbohydrate limited conditions. The observed reduction in Lachnospiraceae, along with decreased taxonomic redundancy and overall diversity in MCD and LCD-fed mice, suggests loss of functional diversity and ecological resilience within the gut microbiota.

Over the past several decades, dietary patterns across many populations have shifted markedly from traditional, fiber-rich whole foods toward diets dominated by processed and ultra-processed products. These modern diets, while often calorically sufficient and fortified with essential micronutrients, tend to be rich in rapidly digestible carbohydrates derived from a limited set of sources, most notably corn-based sweeteners and starches (Gillespie et al., 2023). This reduction in carbohydrate complexity parallels a broader trend toward dietary homogenization, with emerging evidence linking such patterns to diminished gut microbial diversity and increased risk of metabolic, inflammatory, and immune-mediated diseases, including obesity, type 2 diabetes, allergies, and inflammatory bowel disease (Severino et al., 2024). While our study did not focus on a specific health outcome, findings from our mouse model underscore the microbiome’s sensitivity to even modest changes in dietary carbohydrate diversity, independent of total carbohydrate content or caloric load. These alterations, while not accompanied by overt changes in host physiology in the short term, may represent early ecological disruptions that could precede health outcomes dependent on microbiota resilience and functional capacity. As dietary simplification continues to characterize the global food supply, understanding how specific features of diet shape microbial ecosystems will be critical for developing nutritional strategies to maintain or restore gut microbial resilience. Future research should explore the long-term consequences of carbohydrate simplification on microbial functions, integrate more sensitive host physiological measurements, and consider how diet-microbiome interactions influence disease susceptibility.

## FUNDING

This work was supported by the National Science Foundation [NSF-2025541 to AMS and CF] and National Institutes of Health [grant number R35GM150609 to AMS].

## ACKNOWLEDGEMENTS

We would also like to acknowledge Clemson University for generous allotment of compute time on Palmetto cluster, supported by the National Science Foundation under Grant Nos. MRI# 2024205, MRI# 1725573, and CRI# 2010270. We also thank the Clemson University Genomics and Bioinformatics Facility for their help in sample processing and analysis, which receives support from the College of Science and two Institutional Development Awards (IDeA) from the National Institute of General Medical Sciences of the National Institutes of Health (P20GM146584 and P20GM139769).

## Notes

https://github.com/SeekatzLab/RoL-FunctionalRedundancy-microbiota

### Competing Interest Statement

The authors have declared no competing interest.

