## Supplemental Material for "Reduced carbohydrate complexity alters gut microbial structure independent of total carbohydrate intake"

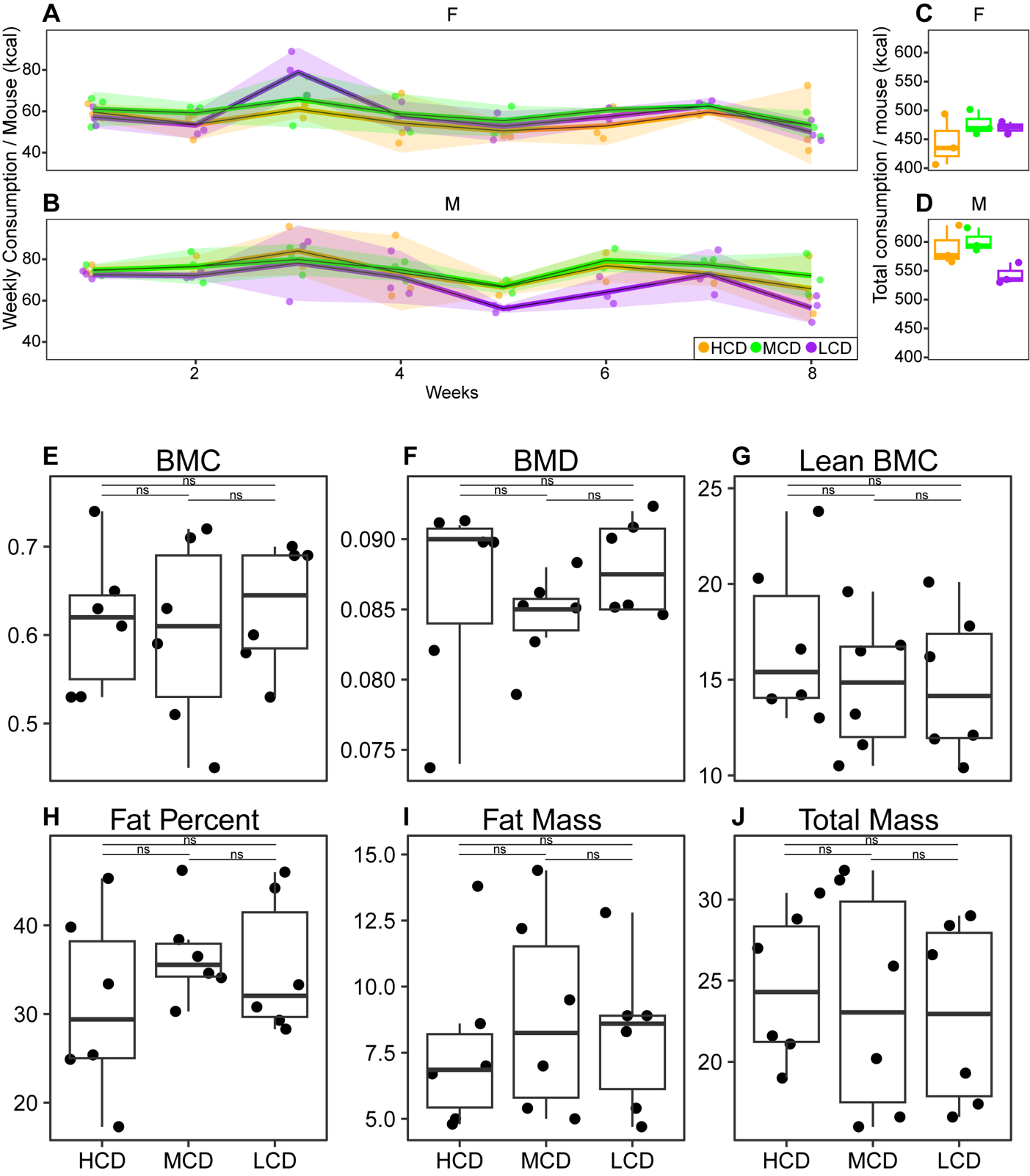

**Figure S1. Dietary carbohydrate complexity did not significantly affect physiological parameters in mice. A-B)** Weekly caloric intake (kcal) of female **A)** and **B)** male mice throughout the study. Total cumulative caloric intake (kcal) by the end of the study for **C)** female and **D)** male mice. **E-J)** DEXA scan-derived physiological measurements: **E)** bone mineral content (BMC), **F)** bone mineral density (BMD), **G)** lean BMC, **H)** body fat percentage, **I)** fat mass, and **J)** total body mass.

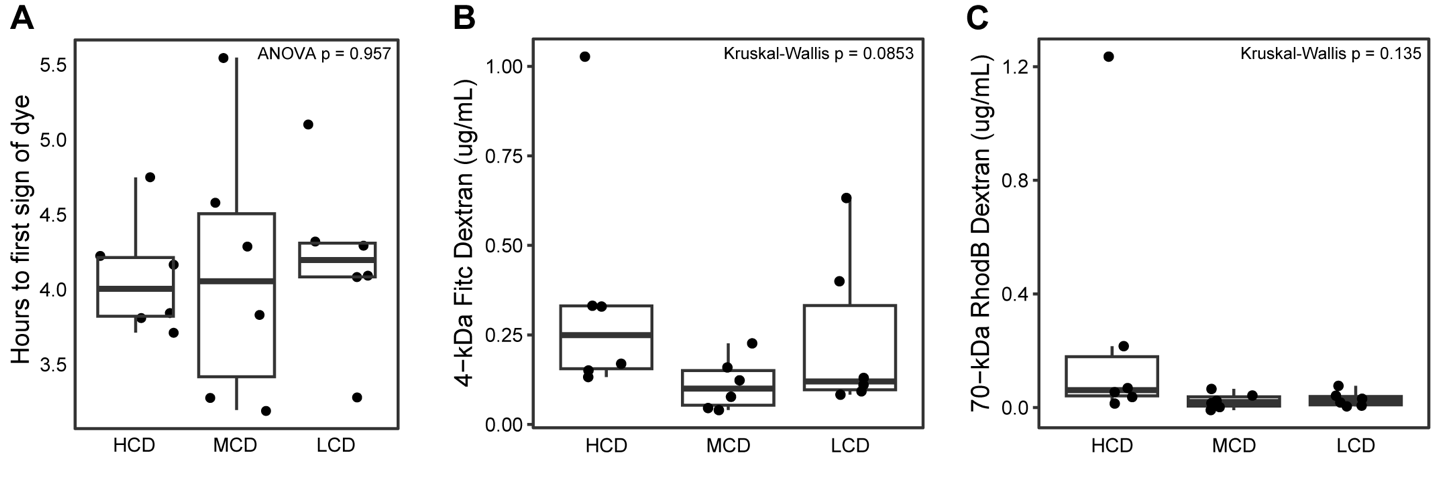

**Figure S2. Dietary carbohydrate complexity did not significantly affect intestinal permeability. A)** Gastrointestinal transit time and **B)** intestinal permeability measured by 4-kDa FITC-dextran and **C)** 70-kDa Rhodamine B concentrations in serum.

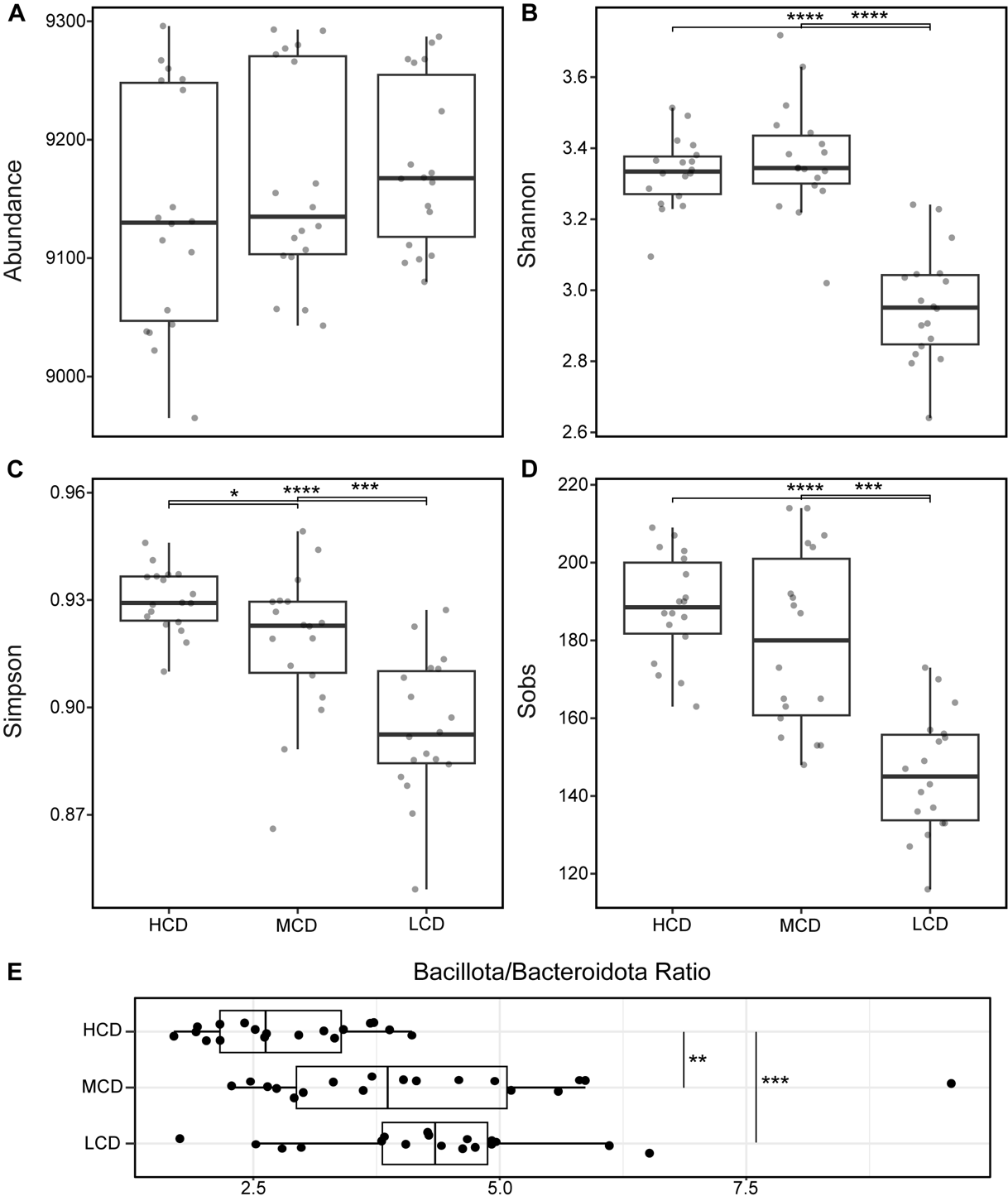

**Figure S3. Lower complexity diets reduce gut microbiota diversity.** Alpha diversity indices based on ASVs: **A)** Abundance **B)** Shannon index **C)** Simpson index and **D)** Observed sequences (Sobs). **E)** Bacillota/Bacteroidota ratio across diet group. Statistical analysis was performed using Kruskal-Wallis tests followed by pairwise Wilcoxon rank-sum tests (*p<0.05, **p<0.001, ***p<0.0001).

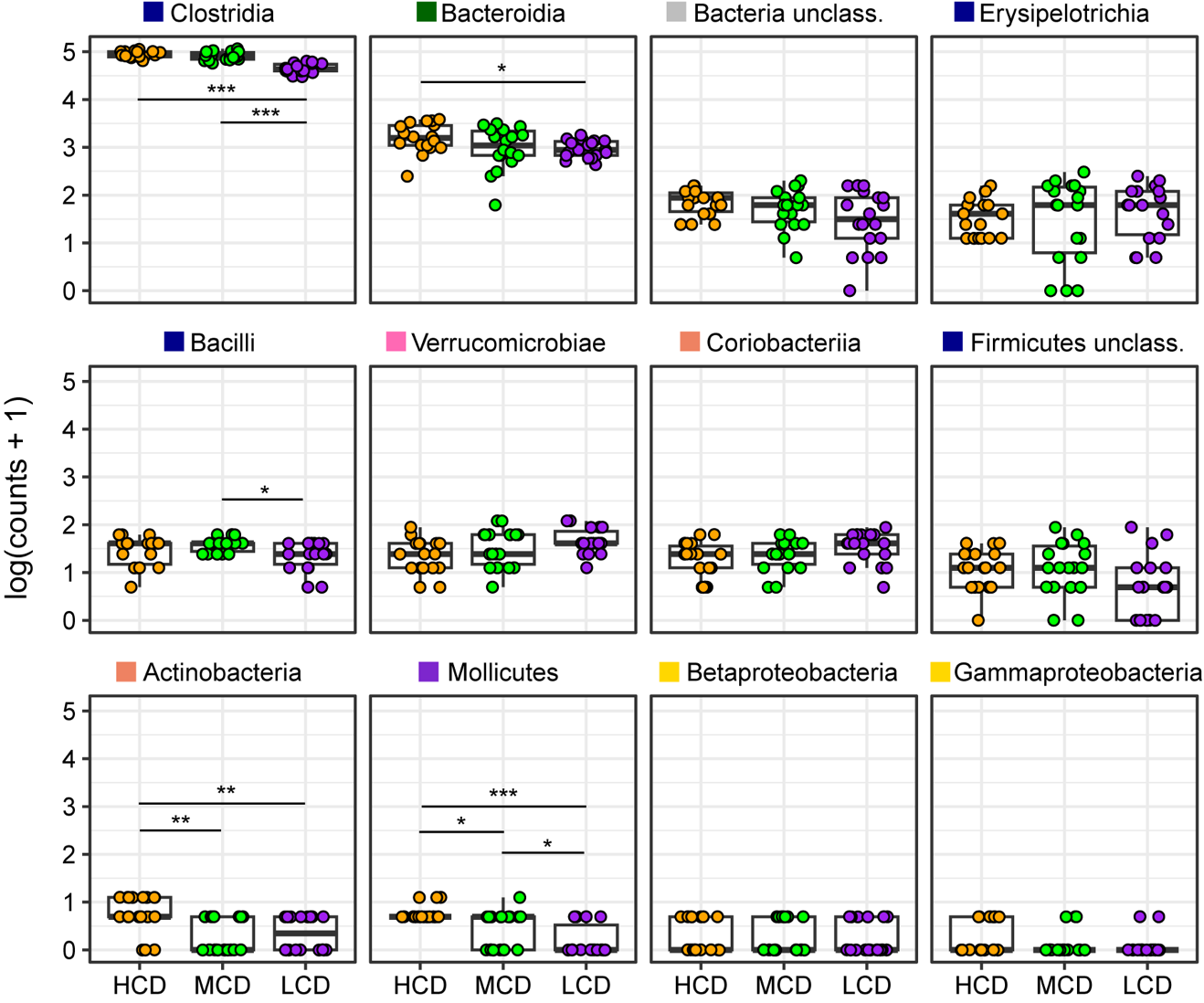

**Figure S4. Taxonomic redundancy at the class level.** Log-transformed (ln[count + 1]) unique ASV counts per mouse at the class level. Statistical comparisons performed using Kruskal-Wallis test with post hoc pairwise Wilcoxon rank-sum tests (*p<0.05, **p<0.001, ***p<0.0001).

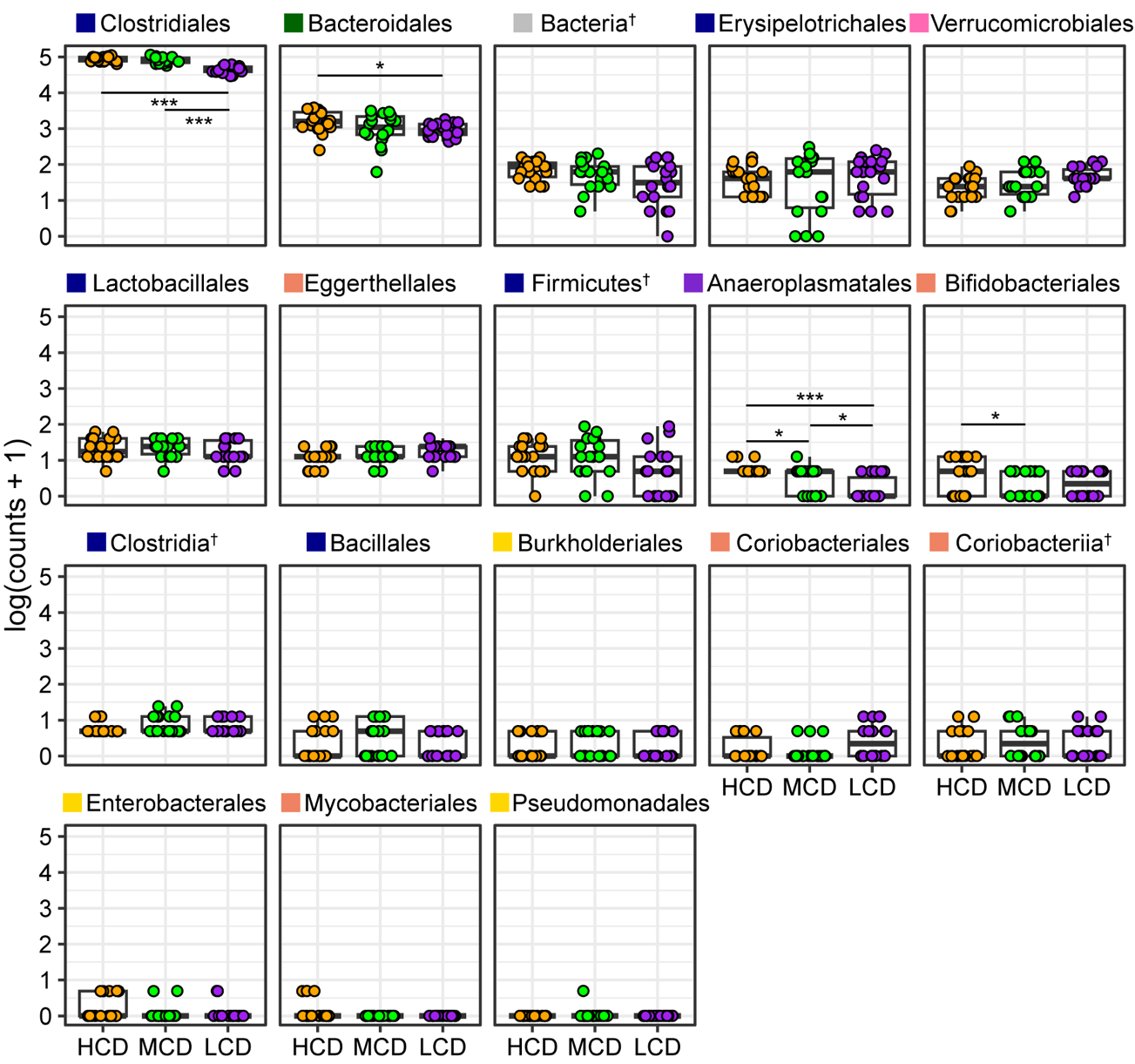

**Figure S5. Taxonomic redundancy at the order level.** Log-transformed (ln[count + 1]) unique ASV counts per mouse at the order level. Statistical comparisons performed using Kruskal-Wallis test with post hoc pairwise Wilcoxon rank-sum tests (*p<0.05, **p<0.001, ***p<0.0001).

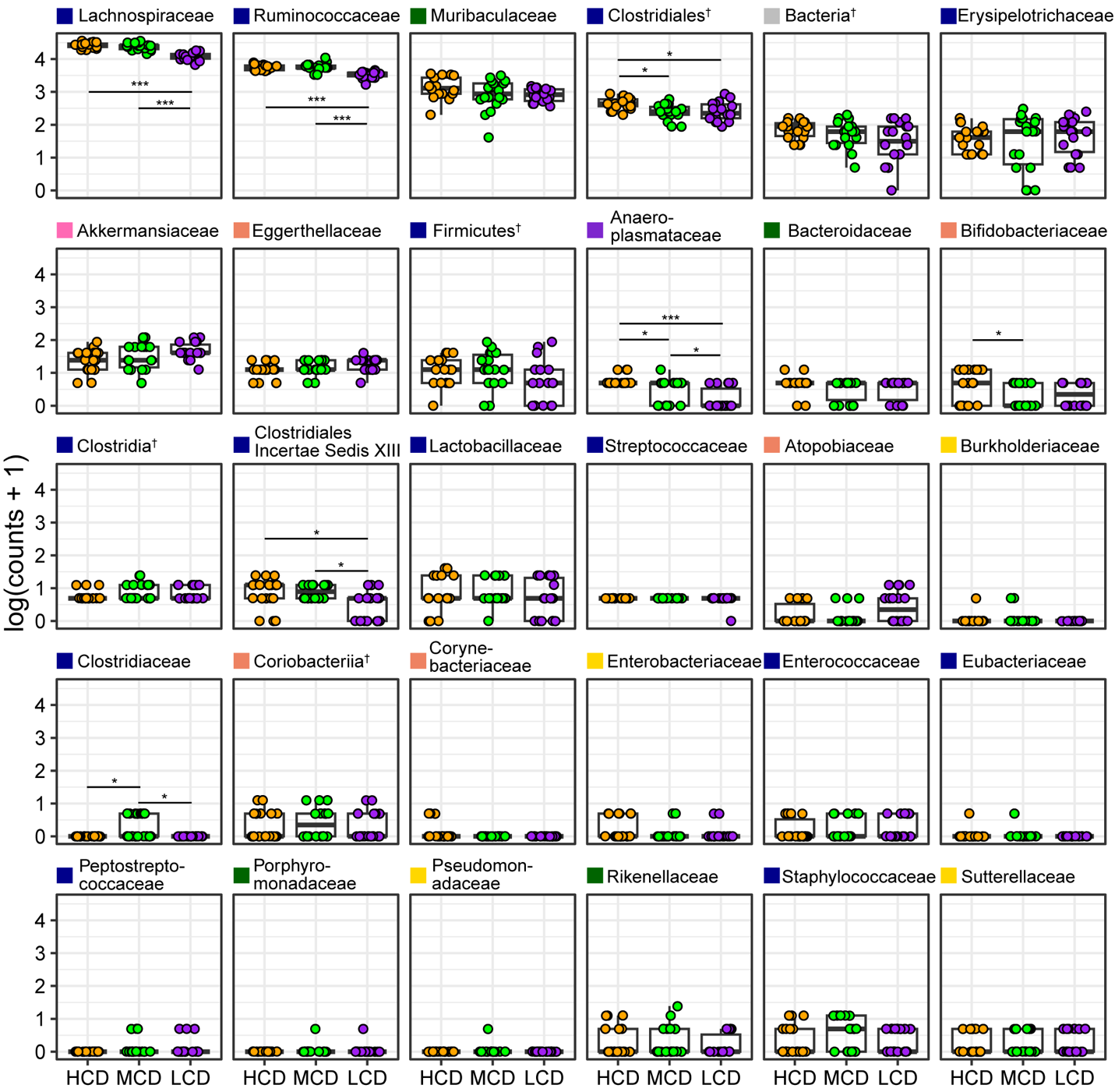

**Figure S6. Taxonomic redundancy at the family level.** Log-transformed (ln[count + 1]) unique ASV counts per mouse at the family level. Statistical comparisons performed using Kruskal-Wallis test with post hoc pairwise Wilcoxon rank-sum tests (*p<0.05, **p<0.001, ***p<0.0001).

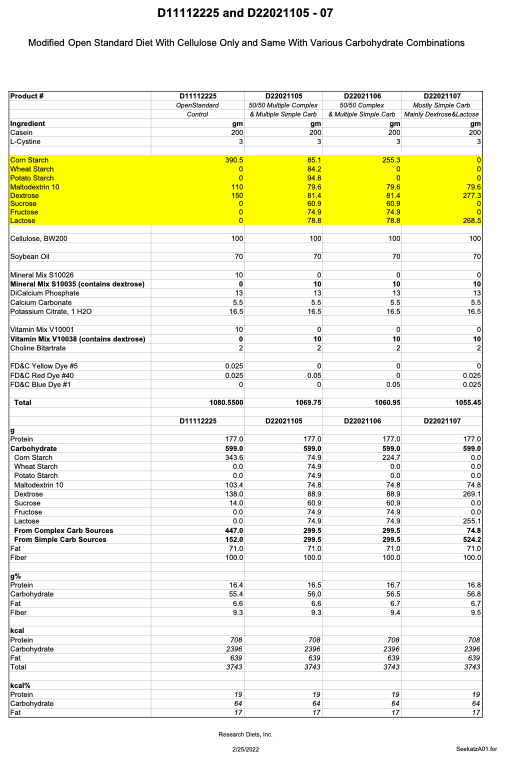
**SUPPLEMENTARY TABLES**

**Table S1. Diet breakdown.** Summary of customized diets compared to Research Diets Open Standard Control (D11112225), High-complexity Diet (HCD, D22021105), Mid-complexity Diet (MCD, D22021106), and Low-complexity Diet (LCD, D22021107).

**Table S2. Overview of project metadata.** Summary of metadata associated with the project, including sample identifiers, and experimental conditions.

| **Genus** | **Phylum** | **HCD vs MCD** | **HCD vs LCD** | **MCD vs LCD** |
| --- | --- | --- | --- | --- |
| *Lachnospiraceae* | Firmicutes | * | *** | *** |
| *Ruminococcaceae* | Firmicutes | n.s | *** | *** |
| *Clostridiales* | Firmicutes | * | * | n.s |
| *Muribaculaceae* | Bacteroides | * | * | n.s |
| *Acetatifactor* | Firmicutes | n.s | * | * |
| *Dysomobacter* | Firmicutes | * | n.s | * |
| *Paramuribaculum* | Bacteroides | n.s | ** | * |
| *Duncaniella* | Bacteroides | n.s | n.s | * |
| *Intestimonas* | Firmicutes | n.s | * | ** |
| *Clostridium XIVa* | Firmicutes | * | n.s | * |
| *Dorea* | Firmicutes | * | n.s | n.s |
| *Ihubacter* | Firmicutes | n.s | * | * |
| *Anaeroplasma* | Tenericutes | * | *** | * |
| *Bifidobacterium* | Actinobacteria | * | n.s | n.s |
| *Eisenbergiella* | Firmicutes | * | n.s. | * |
| *Staphylococcaceae* | Firmicutes | n.s | n.s | ** |
| *Butyricicoccus* | Firmicutes | n.s | * | n.s |
| *Clostridium sensu stricto* | Firmicutes | * | n.s | * |

**Table S3. Pairwise statistical comparisons of taxonomic redundancy at the genus level across diet groups.** Summary of Kruskal-Wallis tests followed by pairwise Wilcoxon rank-sum tests on log_2_(count + 1) unique ASV counts, averaged per diet group for each genus in the dataset, as shown in Figure 4. Significance levels are indicated as *p<0.05, **p<0.001, and ***p<0.0001.
